# Lasting Increases in Neuronal Activity and Serotonergic Receptor Expression Following Gestational Chlorpyrifos Exposure

**DOI:** 10.1101/2025.05.09.653166

**Authors:** Jeffrey A. Koenig, Nathan Cramer, Jimmy Olusakin, Mary Kay Lobo, Asaf Keller

## Abstract

Perinatal exposure to the organophosphorus insecticide chlorpyrifos (CPF) is associated with an increased incidence of neurodevelopmental disorders, such as autism spectrum disorder. While these behavioral detriments have been modeled in rodents, the underlying functional alterations in the developing brain are largely unknown. Previous reports using a rat model have identified alterations to both inhibitory synaptic transmission and serotonergic (5-HT) receptor binding in the cortex following developmental CPF exposure.

Here, we use a rat model of gestational CPF exposure to investigate whether this altered inhibitory activity is driven by increased spontaneous firing of inhibitory interneurons and altered 5-HT receptor expression. Using cell-attached *ex vivo* electrophysiology in young rats of both sexes, we identified a significant increase in the number of spontaneously firing neurons in the somatosensory cortex of CPF-exposed offspring. Analysis of action potential metrics identified a sub-set of these neurons as fast-spiking parvalbumin (PV) interneurons. Immunohistochemical labeling of c-Fos, a marker of neuronal activity, further revealed a pronounced increase in activity of neurons of the somatosensory cortex in both juvenile and adult rats that had been gestationally exposed to CPF. Finally, RNAscope *in situ* hybridization showed an increase in the expression of the inhibitory receptor 5-HT_1B_ in PV neurons. The data here demonstrate that gestational exposure to CPF results in persistent hyper-excitation of the somatosensory cortex, potentially through increased expression of the receptor 5-HT_1B_ and resulting disinhibition. These neurophysiological effects may contribute to the established behavioral outcomes resulting from gestational exposure to CPF and offer guidance for novel preventative interventions.

**SIGNIFICANCE STATEMENT:** We report persistent increases in spontaneous neuronal firing in the somatosensory cortex following a brief gestational exposure to the organophosphorus insecticide CPF in rats. This occurred in conjunction with increased expression of the 5-HT_1B_ receptor measured specifically in PV interneurons. The hyper-excitability of the somatosensory cortex described here agrees with the established hyper-sensitivity to sensory stimulation seen in neurodevelopmental disorders such as autism spectrum disorder. These results offer a possible mechanistic framework underlying the neurophysiological effects associated with early life exposures to the organophosphorus insecticide CPF.

## INTRODUCTION

Chlorpyrifos (CPF) is an organophosphorus insecticide ubiquitously used in agriculture. Like other organophosphorus compounds, CPF is toxic to non-target species, including humans, primarily through potent inhibition of the enzyme acetylcholinesterase (Taylor, 2011). The acute cholinergic effects of CPF exposure have long been understood, but more recent attention has been given to sub-acute environmental exposures that are not associated with overt signs and symptoms of acute toxicity or significant inhibition of acetylcholinesterase. Several longitudinal studies have linked these exposures, especially during pregnancy, to an increased incidence of neurodevelopmental disorders such as attention deficit/hyperactivity disorder and autism spectrum disorder (Rauh et al., 2012; Schmidt et al., 2017; Shelton et al., 2014). While this has led to more restrictions on the application of CPF, its worldwide use continues to grow (ECHA, 2023).

The developmental neurobehavioral effects of CPF have been modeled in several animal species, including rats, mice, and guinea pigs. Perinatal exposures of these animals to doses of CPF that do not induce acute toxicity results in increased anxiety-related behaviors, learning and working memory deficits, and alterations to somatosensory function (Aldridge et al., 2005; Levin et al., 2001; Mamczarz et al., 2016; Muller et al., 2014). Investigations into non-cholinergic neurochemical alterations have primarily implicated serotonergic (5-HT) signaling systems. Following perinatal CPF exposure in rats, binding of 5-HT_1A_ and 5-HT_2_ receptors is increased at least until 5 months of age (Aldridge et al., 2004, 2003; Slotkin and Seidler, 2005). However, 5-HT receptors are diversely expressed, both pre and postsynaptically, and on both inhibitory and excitatory neurons (Celada et al., 2013; Palacios, 2016), making it difficult to predict the functional consequences of this increase in receptor binding. Further, whether these changes in receptor binding reflect changes in receptor expression or in receptor affinity remains unknown.

Following sub-acute gestational exposure to CPF in rats, there are persistent alterations to inhibitory synaptic signaling and use-dependent plasticity in rat somatosensory (barrel) cortex (Koenig et al., 2025). We hypothesize that these anomalies are driven by disinhibition and increased activity of parvalbumin (PV) neurons. PV neurons are a unique class of fast-spiking inhibitory neurons that provide dense input and inhibition to neighboring cells (Packer and Yuste, 2011). In addition to serving as a primary inhibitory input to pyramidal neurons and to other PV neurons, they are critical for the proper refinement of use dependent plasticity (Rupert and Shea, 2022; Trachtenberg, 2015; Vickers et al., 2018). Dysregulation of PV neuron activity has been implicated in several neurological disorders, including autism spectrum disorder (Filice et al., 2020; Marín, 2012). PV neurons are particularly sensitive to toxin exposure during development (Dendrinos et al., 2011; Reid et al., 2021; Stansfield et al., 2015).

In the present study, we test the hypothesis that changes in cortical functions after gestational exposure to CPF result in lasting *in vivo* increases in neuronal activity and in cell-specific changes in 5-HT receptor expression. We focus on the somatosensory (barrel) cortex, as its development is highly characterized (Erzurumlu and Gaspar, 2012; Yang et al., 2018), and because both PV neurons and proper 5-HT signaling play a key role in its maturation and function (Kimura and Itami, 2019; Miceli et al., 2017, 2013; Nowicka et al., 2009; Sachidhanandam et al., 2016; Teissier et al., 2017).

## MATERIALS AND METHODS

### Animals and treatment

All procedures adhered to the Guide for the Care and Use of Laboratory Animals and approved by the Institutional Animal Care and Use Committee at the [Author Institution]. Male and female Long-Evans rats were acquired from Charles River Laboratories and bred in our vivarium. Animals were housed under a standard 12/12 hour light/dark cycle (0600 on/1800 off) and had *ad libitum* access to food and water. Pregnancy was confirmed by the presence of sperm in a vaginal lavage collection and set as gestational day (GD) 0. CPF (Chem Service, West Chester, PA) was dissolved in a 50/50 mixture of DMSO/peanut oil and injected subcutaneously (0.5 mL/kg) once daily on GD 18-21 at a dose of 5 mg/kg.

Control dams received vehicle (DMSO/peanut oil) injections on the same schedule. Litters were culled to a maximum size of 10 by PND 3. Dams were pair-housed until GD 18. Both male and female progeny were used in all experiments, unless noted otherwise in Results.

### Slice electrophysiology

As previously described (Alipio et al., 2021), animals were deeply anesthetized with ketamine/xylazine, the brains were removed, and 300 μm coronal slices containing the primary somatosensory cortex were prepared. Slices were placed in a recording chamber continuously perfused (1.5 mL/min) with carbogen saturated artificial cerebrospinal fluid (ACSF) containing: 124 mM sodium chloride, 2.5 mM potassium chloride, 1.25 mM monosodium phosphate, 24 mM sodium bicarbonate, 12.5 mM glucose, 2 mM magnesium sulfate heptahydrate, and 2 mM calcium chloride dihydrate (Sigma-Aldrich). The recording pipette (4-6 MΩ) was filled with ACSF solution. Inhibitory interneurons in the superficial layers (L 2/3 and L 4) of the barrel cortex were targeted based on morphology and lack of prominent apical dendrite. Cell-attached recordings were performed in voltage clamp mode (0 pA holding current) with a loose seal resistance (< 200 MΩ) (Perkins, 2006). Seal resistance was measured before and after recording to ensure patch stability. A cell was considered “active” if there were any spontaneous fast-current transients representing action potentials within an ∼3-minute recording period.

### Histology

c-Fos expression was visualized through immunohistochemistry. Both juvenile (PND 12-13) and adult (> PND 90) animals were used in this experiment. Juvenile animals were removed from their littermates and dam before being quickly (<15 minutes) euthanized. Adult animals were monitored in their home cage, absent enrichments, for 2 hours prior to collection, as previously described (Filipkowski et al., 2000). There was no discernible, qualitative difference in the mobility or other activity between animals from CPF-and vehicle-exposed groups. Animals were deeply anesthetized by intraperitoneal injection of ketamine/xylazine (80 and 10 mg/kg) and transcardially perfused with ice-cold PBS followed by 10% neutral buffered formalin (NBF). The brains were extracted and incubated in NBF at 4°C for 24 hours. Brains were then cryoprotected with increasing concentrations of sucrose (10, 20, and 30%) in PBS, allowing the brain to sink at each concentration (∼24 hours). Following the final cryoprotection step, the brains were embedded in optimal cutting media (OCT) and rapidly frozen. Tissue was stored at-80°C until processing. For juvenile tissue, 12 µm thick sections were cut with a cryostat and mounted on charged slides. Slides were baked at 57°C for 20 minutes then rinsed with PBS for 10 minutes to remove OCT. A hydrophobic barrier was drawn surrounding the tissue. Sections were incubated with blocking buffer (10% normal donkey serum, 0.5% Triton X-100) for 2 hours at room temperature. Blocking buffer was removed and sections were incubated with phospho-c-Fos (Ser32) primary antibody (1:1000, D82C12, Cell Signaling Technology) at 4°C for 24 hours. Slides were rinsed 3 times for 10 minutes in PBS prior to a 1 hour incubation with secondary antibody, donkey anti-rabbit Alexa Fluor Plus 488 (1:500, A32790, Thermo Fisher). Slides were rinsed 3 times for 10 minutes in PBS and mounted with ProLong Gold Antifade Mountant with DAPI (Thermo Fisher). For adult tissue, 40 µm thick sections were cut with a cryostat and transferred to PBS on 12 well plates. Sections were incubated in blocking buffer for 2 hours at room temperature prior to 48 hour incubation in phospho-c-Fos (Ser32) primary antibody (1:500, D82C12, Cell Signaling Technology) at 4°C for 48 hours. Secondary antibody incubation (1:500, donkey anti-rabbit Alexa Fluor Plus 488) was for 2 hours at room temperature prior to mounting. Images were acquired with a Leica Mica microscope at 20x. Analysis was performed with Imaris 10 (Oxford Instruments), quantifying the number of c-Fos positive nuclei within a 500 µm wide column through all layers of the barrel cortex. Only female animals were used in this experiment.

### RNAscope

To measure 5-HT_1B_ receptor expression specifically in PV neurons, RNAscope *in situ* hybridization was performed in conjunction with immunohistochemistry on 12 µm thick sections from fixed-frozen tissue according to the manufacturer supplied protocol (ACD, MK 51-150). Following antigen retrieval, sections were incubated overnight at 4°C with anti-PV antibody (1:2500, PA1-933, Thermo Fisher). RNAscope hybridization was performed using probe 5-hydroxytryptamine receptor 1B (Htr1b, 420361, ACD) and fluorophore Opal 650 (1:1000, Akoya Biosciences). There was a 30 minute incubation with secondary antibody donkey anti-rabbit Alexa Fluor Plus 488 (1:250, A32790, Thermo Fisher) prior to mounting with ProLong Gold Antifade Mountant with DAPI (Thermo Fisher). Images of the barrel cortex were acquired with a Leica SP8 confocal microscope at 40x. Quantification of 5-HT_1B_ transcript puncta co-localized with PV immunohistochemistry labeling was performed using Imaris 10 (Oxford Instruments).

### RNA isolation and RT-qPCR

Tissue punches were collected from the barrel cortex of PND 12-14 rats bilaterally with a 15-gauge punch and stored at-80°C until processing. RNA was extracted and isolated using TRIzol reagent (Thermo Fisher) and the MicroElute total RNA kit (Omega Bio-tek, Inc) according to the manufacturer instructions. RNA concentrations were measured on a NanoDrop spectrophotometer, and 400 ng of cDNA was synthesized using an iScript cDNA synthesis kit (Bio-Rad Laboratories). mRNA expression was measured by RT-qPCR with Perfecta SYBR Green FastMix (Quanta) and a CFX384 system (Bio-Rad Laboratories) using the ΔΔCT method. GAPDH was used as the housekeeping gene. The primer sequences were: 5-HT_1B_ receptor (Htr1b, NM_022225.3) forward, 5′-AGAAGAAACTCATGGCCGCT and reverse, 5′-GGGGAGCCAGCACACAATAA; GAPDH (Gapdh, NM_017008.4) forward, 5′-TGGCCTCCAAGGAGTAAGAA and reverse, 5′-TGTGAGGGAGATGCTCAGTG.

### Statistical analysis

Statistical analyses were conducted with GraphPad Prism 10 (Boston, MA). Statistical significance was set as *p* < 0.05. Parametric tests were used if the appropriate assumptions were met. Otherwise, nonparametric tests were used. The statistical tests used are listed in each figure legend. There were no differences between rats of different litters in the reported endpoints, thus they were combined for analysis. Sex differences were investigated where powered and noted in Results. In all experiments we adhere to accepted standards for rigorous study design and reporting to maximize the reproducibility and translational potential of our findings, as described in Landis et al (2012) and in ARRIVE (Animal Research: Reporting In Vivo Experiments) Guidelines. We performed a power analysis to estimate the minimal sample size for each experiment, using G*Power software (Faul et al., 2007).

## RESULTS

### Spontaneous neuronal activity

Gestational CPF exposure results in a lasting increase in inhibitory inputs to pyramidal neurons (Koenig et al., 2025). To determine if this is driven by increased spontaneous activity of inhibitory neurons, we employed a cell-attached voltage clamp technique in a brain slice preparation from juvenile (PND 12-20) rats. Interneurons were targeted based on their morphology in the superficial layers (layers 2/3 and layer 4) of the primary somatosensory (barrel) cortex. Example cell-attached traces from a vehicle and CPF treated animal are seen in Figure 1A. Action potential currents can be seen in the representative trace from a CPF exposed animal; these are absent in the neuron from the vehicle treated animal. There was a greater proportion of neurons displaying spontaneous activity (Fig 1B) in animals that had gestational CPF exposure (20 of 28, 71%) compared to neurons from vehicle-treated animals (1 of 25, 4%). These findings demonstrate that gestational CPF exposure results in a persistent increase in spontaneous neuronal activity of suspected inhibitory neurons.

**Figure 01.**
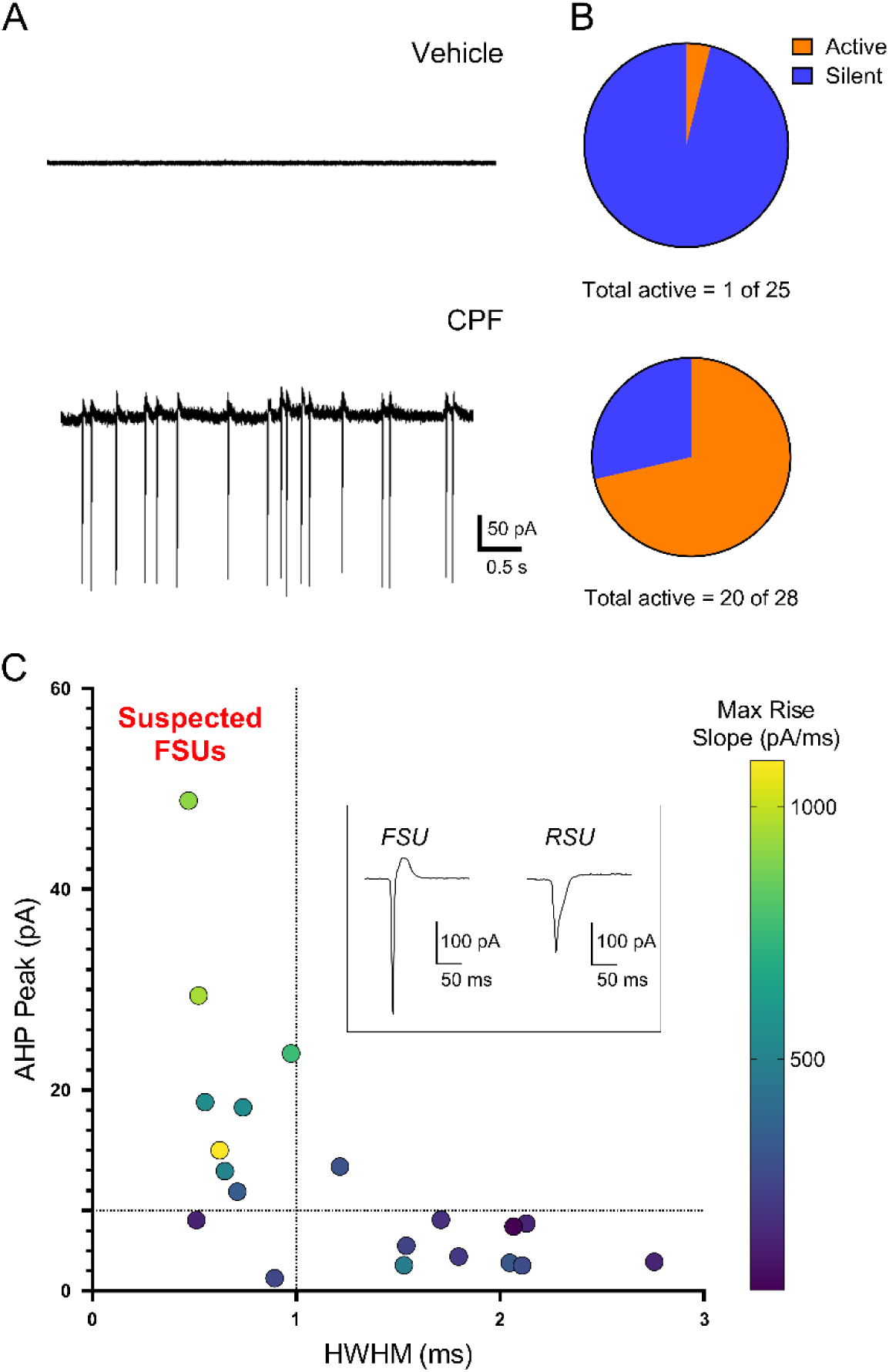
Gestational chlorpyrifos exposure increases spontaneous activity of interneurons in the somatosensory (barrel) cortex. **A.** Example cell-attached recording traces of suspected interneurons from vehicle and CPF exposed animals. **B.** A greater proportion of neurons had spontaneous activity (p < 0.0001, Fisher’s exact test) in the treated (CPF) group (20 of 28 neurons from 3 animals [2 male, 1 female]) compared to vehicle (1 of 25 neurons from 3 animals [1 male, 2 female]). **C.** A subset of spontaneously active neurons (8 of 20) were identified as suspected fast-spiking units (FSUs) based on their after-hyperpolarization peak (AHP) being > 8 pA and having an action potential half-width (HWHM) < 1 ms. Inset shows example action potential traces from a FSU and a regular-spiking unit (RSU).

We used previously established metrics to distinguish PV fast spiking units (FSUs) from regular spiking units (RSUs) (Connors et al., 1982; McCormick et al., 1985; Murray and Keller, 2011). Fig 1C (inset) shows an example FSU with a characteristic short action potential duration and prominent afterhyperpolarization (AHP), absent from the RSU.

Neurons with both an AHP peak > 8 pA and a half width at half maximum (HWHM) < 1 ms were designated as suspected FSUs. Of the 20 spontaneously active neurons in the CPF group, 8 were characterized as FSUs and 12 as RSUs (Fig 1C). Those identified as RSUs could comprise pyramidal neurons or other classes of interneurons, such as somatostatin or serotonin receptor 5HT_3A_ expressing (Rudy et al., 2011).

### c-Fos expression

As a second metric for increased neuronal activity, we evaluated increases in the expression of phosphorylated c-Fos, a product of an immediate early gene that serves as a reliable marker of neuronal activity, and that is quickly upregulated following neuronal activation (Dragunow and Faull, 1989; Morgan et al., 1987). In both juvenile and adult vehicle-exposed offspring, labeling of c-Fos positive nuclei was sparse across all layers (Fig 2A,B). In contrast, in the CPF-exposed offspring there was a marked increase in c-Fos labeling relative to vehicle-treated. In the juvenile age group, the numbers of c-Fos-positive nuclei were 8-fold higher in layers 2/3, 18-fold higher in layer 4, but remained the same in layers 5/6 of the barrel cortex of offspring exposed to CPF compared to vehicle. In the adult age group, numbers of c-Fos positive nuclei were 6-fold higher in layers 2/3, 16-fold higher in layer 4, and 4-fold higher in layers 5/6 of the barrel cortex of offspring exposed to CPF compared to vehicle (Fig 2B). These data further support the hypothesis that CPF exposure results in persistent enhanced activation of cortical neurons.

**Figure 02.**
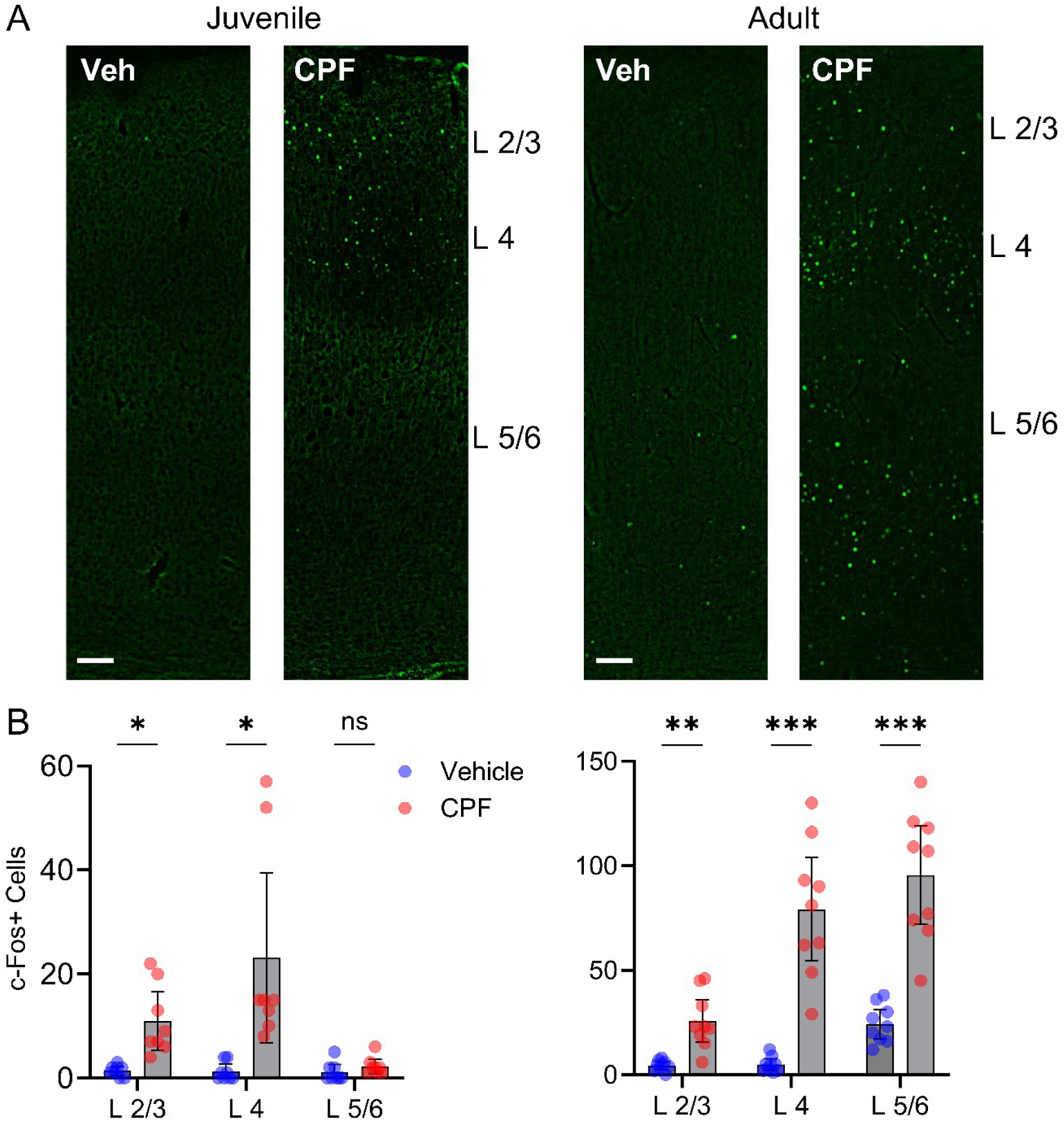
Gestational chlorpyrifos exposure increases c-Fos expression in both juvenile and adult animals. **A.** Representative immunohistochemistry images of the barrel cortex of juvenile (PND 12-13) and adult (> PND 90) female animals showing c-Fos positive nuclei. Scale bar 100 µm. **B.** The number of c-Fos positive nuclei is higher in CPF exposed juvenile animals (n = 9 sections from 3 animals) in layers 2/3 (t(7.4) = 4.0, p = 0.005) and layer 4 (t(7.1) = 3.1, p = 0.03), but not layers 5/6 (t(13.9) = 1.3, p = 0.22). In adult animals (n = 9 sections from 3 animals), that number is higher in layers 2/3 (t(8.6) = 4.7, p = 0.001), layer 4 (t(8.2) = 6.9, p < 0.001), and layers 5/6 (t(9.4) = 6.7, p < 0.001, Welch’s unpaired t test). Mean ± 95% CI (**B**).

### 5-HT_1B_ receptor expression

A potential mechanism driving both the increased spontaneous activity of neurons seen here, and in previously reported enhancement of inhibitory synaptic transmission (Koenig et al., 2025) is presynaptic disinhibition of PV neurons. As stated above, there is a persistent elevation of binding to 5-HT_1A_ and 5-HT_2_ receptors in the brain of rats following gestational CPF exposure (Aldridge et al., 2004, 2003). While the 5-HT_1A_ receptor is both inhibitory and presynaptically expressed, it functions primarily as an autoreceptor on 5-HT neurons (Altieri et al., 2013; Verge et al., 1985). The 5-HT_1B_ receptor acts presynaptically to suppress transmitter release and is highly expressed on inhibitory neurons, potentially driving this disinhibition (Bramley et al., 2005; Egeland et al., 2011). Increased activity of 5-HT_1B_ receptors on inhibitory neurons could result in the disinhibition of interneurons that receive synaptic inputs form those inhibitory neurons. Lasting changes in the expression of 5-HT_1B_ receptors after prenatal CPF exposure have yet to be studied.

Here, we employed RNAscope *in situ* hybridization to label 5-HT_1B_ transcripts, in conjunction with immunohistochemistry labeling of PV neurons in the barrel cortex. Fig 3A shows a representative image of immunohistochemistry-labeled PV neurons (green) co-localized with 5-HT_1B_ transcripts (red puncta). Each puncta corresponds to a single 5-HT_1B_ mRNA transcript (Wang et al., 2012). Quantification of puncta specifically co-localized with PV labeling (Fig 3B) revealed 65% higher number of transcripts per cell in male offspring exposed to CPF compared to vehicle (veh: 2.6 ± 0.46; CPF: 4.3 ± 0.50) and 9% greater number of transcripts per cell in female offspring exposed to CPF compared to vehicle (veh: 8.7 ± 0.61; CPF: 9.5 ± 0.51). When comparing by sex, vehicle-exposed females had a 236% greater number of 5-HT_1B_ transcripts per PV neuron than vehicle-exposed males, and CPF-exposed females had a 122% greater number of transcripts per PV neurons than CPF-exposed males. There was not a significant interaction between sex and treatment (p = 0.24).

**Figure 03.**
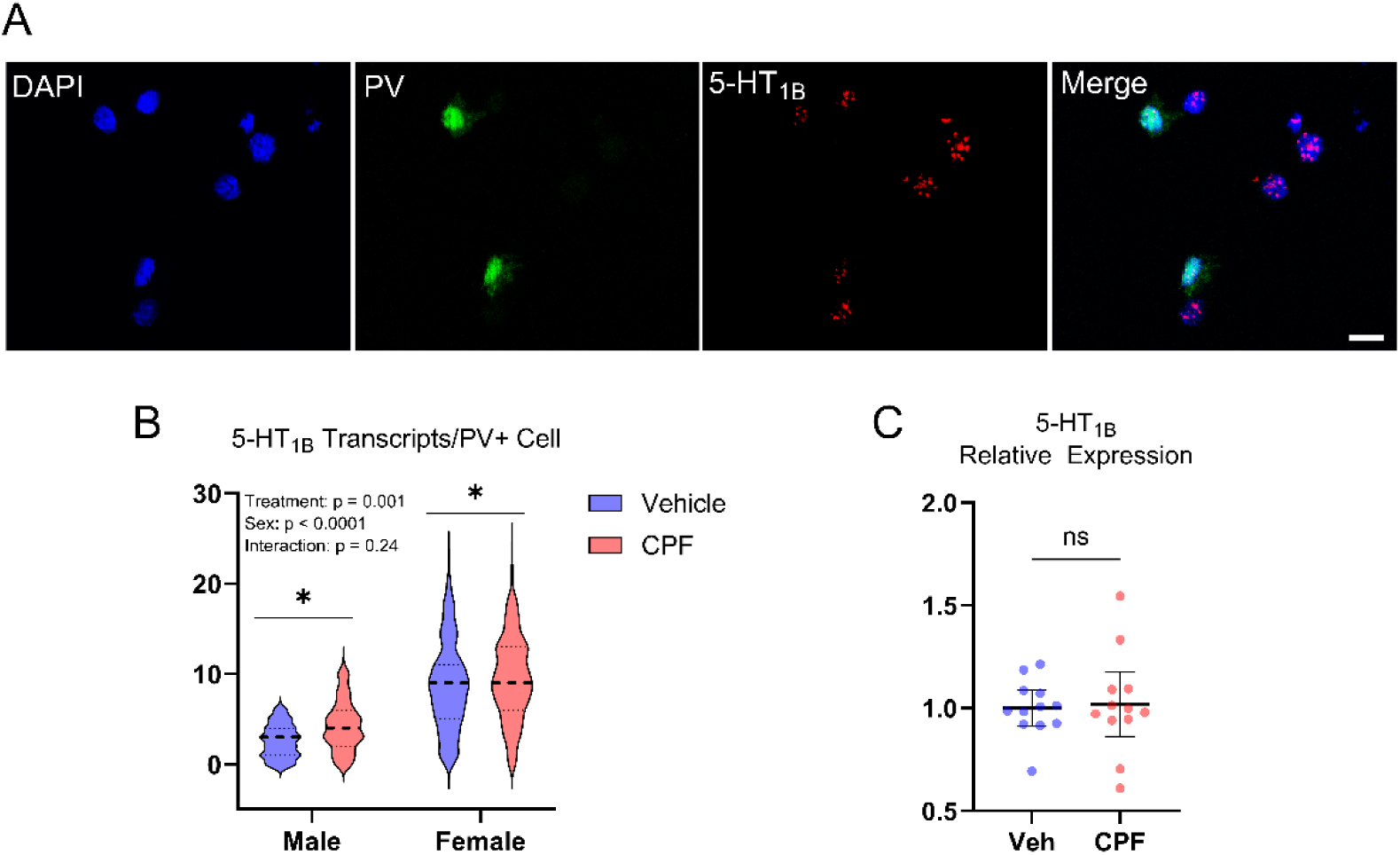
Gestational chlorpyrifos exposure increases expression of the 5-HT_1B_ receptor in PV neurons. **A.** Representative images showing immunohistochemistry labeling of PV interneurons and *in situ* hybridization RNAscope labeling of 5-HT_1B_ receptor transcripts in the barrel cortex. Scale bar is 10 µm. **B.** Multiple comparison analysis demonstrates higher 5-HT_1B_ receptor expression following CPF exposure in both males (p = 0.03, vehicle n = 62 cells from 2 animals, CPF n = 116 cells from 2 animals) and females (p = 0.03, vehicle n = 251 cells from 3 animals, CPF n = 316 cells from 3 animals, Holm-Šídák’s test) compared to vehicle. There is a significant effect for both treatment (F(1,740) = 10.36, p = 0.001) and sex (F(1,740) = 219.8, p < 0.0001), but not for the interaction of the two (F(1,740) = 1.357, p = 0.24, two-way ANOVA). **C.** 5-HT_1B_ receptor expression in the barrel cortex measured by RT-qPCR did not demonstrate a difference (t(17.2) = 0.23, p = 0.82, Welch’s unpaired t test) between vehicle (n = 12 animals [7 male, 5 female]) and CPF (n = 12 animals [9 male, 3 female]) groups. Violin plot with median and quartiles (**B**). Mean ± 95% CI (**C**).

Measurement of total 5-HT_1B_ expression in the barrel cortex using RT-qPCR revealed no difference in the relative expression between vehicle (1.0 ± 0.07) and CPF exposed (1.0 ± 0.16) groups (Fig 3C). The lack of difference seen in these results could be attributed to the non-specific nature of the expression data. If changes in 5-HT_1B_ expression were limited to specific cell types, such as PV neurons, this global measurement could effectively washout any difference. Overall, these findings support the role of increased expression of the 5-HT_1B_ receptor, particularly in inhibitory PV neurons, in driving spontaneous neuronal activity following CPF exposure.

## DISCUSSION

We hypothesized that gestational exposure to a sub-acute dose of CPF would result in lasting enhancement of spontaneous neuronal activity and an increased expression of the 5-HT_1B_ receptor, particularly in inhibitory neurons. Consistent with these predictions, there was a marked increase in the number of spontaneously active neurons in the barrel cortex, in conjunction with increased expression of c-Fos protein, a marker of neuronal activation. *In situ* hybridization labeling demonstrated an increase in 5-HT_1B_ expression in PV neurons following CPF exposure.

### Spontaneous neuronal activity

Altered inhibitory synaptic transmission following a sub-acute gestational exposure to CPF has been previously reported (Koenig et al., 2025). This presented as an increase in spontaneous inhibitory synaptic events, but also as a reduction in presynaptic release probability of GABA onto layer 2/3 excitatory pyramidal neurons. These phenomena could be explained by the disinhibition and increase in spontaneous firing of PV neurons, the primary inhibitory input to pyramidal neurons (Packer and Yuste, 2011; Yang et al., 2016).

Here, we demonstrate this increase in spontaneous firing through cell-attached electrophysiology: 71% of the neurons recorded in CPF exposed animals showed spontaneous firing, whereas only 4% of those in the control group fired spontaneously (Fig. 1B).

Only a subset of the spontaneously firing neurons (40%) were identified as PV fast-spiking neurons, based on their action potential metrics (Connors et al., 1982; McCormick et al., 1985). The remaining were identified as regular spiking units (RSUs), and these may include excitatory neurons, as well as other classes of inhibitory cells, such as somatostatin expressing neurons (Rudy et al., 2011). Somatostatin neurons, like their PV counterparts, provide inhibitory input to layer 2/3 pyramidal neurons (Adesnik et al., 2012; Xu et al., 2013), and an increase in their spontaneous firing may contribute to the increase in inhibitory inputs reported previously (Koenig et al., 2025).

PV neurons reach anatomical and functional maturity only in late adolescence, and their maturation is critically sensitive to use-dependent changes in activity (Dehorter et al., 2015; Miller et al., 2011). Dysregulation of PV neuron activity is causally linked to a number of neuropsychiatric disorders, such as schizophrenia, autism spectrum disorder, substance use disorders, and depression (Ferguson and Gao, 2018; Filice et al., 2020; Nahar et al., 2021; Selten et al., 2018). Imbalance in excitatory/inhibitory synaptic transmission has also been implicated in these pathologies (Ferranti et al., 2024; Gao and Penzes, 2015). The hyper-excitability reported here could underly the persistent neurobehavioral detriments associated with early life CPF exposure.

### c-Fos expression

As an additional metric to evaluate enhanced neuronal excitability, we measured c-Fos expression in the barrel cortex of juvenile and adult animals. In agreement with previous reports (Chu et al., 2013), there was only low c-Fos expression in vehicle treated animals. The marked increase in c-Fos expression in CPF exposed animals reported here appeared similar to animals who have undergone robust vibrissae stimulation (Filipkowski et al., 2000).

In juvenile animals that had gestational exposure to CPF, c-Fos labeling revealed neuronal activation in layers 2/3 and 4, but not in deep layers 5/6. This suggests enhanced activity related to thalamocortical activation in layers 2/3 and 4, but not in the summation and output of layers 5/6 (Keller, 1995; Laaris et al., 2000). During the second week of postnatal development, the age of juvenile animals in the present study, layer 2/3 neurons receive more excitatory drive, compared to layer 5 neurons (Kroon et al., 2019). Further, synapses onto layer 2/3 neurons develop earlier than synapses from layer 3 to layer 5 (Clement et al., 2013). Both phenomena could account for the absence of c-Fos labeling in these deep layers of juvenile animals. The robust level of c-Fos expression in adult animals, more than 90 days following the brief gestational exposure, also supports the persistent nature of these CPF-induced neurophysiological alterations.

The cell-types of the c-Fos positive cells were not identified in this experiment. Previous reports have shown that both inhibitory and excitatory cells increase c-Fos expression when exposed to a novel environment (Staiger et al., 2002). An increase in c-Fos expression in PV or other inhibitory neurons is consistent with our electrophysiology results, showing enhanced inhibitory synaptic activity in barrel cortex after gestational CPF exposure (Koenig et al., 2025). However, it is also possible that amplified inhibitory inputs result in paradoxical hyperexcitation of excitatory, pyramidal neurons, as a result of post-inhibitory rebound spiking (Moore et al., 2018; Morishima et al., 2017). As pyramidal and PV neurons form reciprocal connections, this could lead to abnormal response properties of excitatory cortical neurons, and affect cortical oscillations that facilitate cognition and attention (Kim et al., 2016; Sohal et al., 2009; Tan et al., 2019).

Hyper-sensitivity to sensory stimulation is a common phenotype and hallmark in individuals with autism spectrum disorder (Markram and Markram, 2010). Animal models of Fragile X syndrome and valproic acid-induced autism phenotypes have consistently demonstrated persistent hyper-excitability in the somatosensory cortex (Deng and Lei, 2008; Markram and Markram, 2010). The enhanced neuronal spontaneous activity and increased basal c-Fos expression reported here both support this phenomenon and potentially underly the increased incidence of these neurobehavioral detriments seen in populations exposed gestationally to CPF.

### 5-HT_1B_ receptor expression

We have previously reported a reduction in the presynaptic release probability of GABA in CPF exposed animals. We hypothesized this is driven by increased expression of the inhibitory receptor 5-HT_1B_, specifically on PV neurons. As the primary inhibitory input onto PV neurons are other PV neurons (Kubota et al., 2016; Yang et al., 2016), presynaptic disinhibition could lead to their overall increased activity described above. Our *in situ* RNAscope results support this, demonstrating increased 5-HT_1B_ expression in PV neurons of our CPF exposed animals.

The 5-HT_1B_ receptor also plays a critical role in the proper patterning of the rodent barrel cortex (Cases et al., 1996; Rebsam et al., 2002). This altered 5-HT_1B_ expression induced by CPF exposure could be a mechanism driving the sensory processing disorders associated with early OP exposure (Cartier et al., 2018; Silver et al., 2018). It also supports prior findings of alterations to barrel field patterning using this same CPF exposure model (Koenig et al., 2025). Whereas previous studies on the lasting effects of gestational CPF exposure (Aldridge et al., 2004, 2003) only reported changes in global 5-HT receptor activity in particular brain regions, the PV neuron-specific alterations reported here provide mechanistic framework to allow for further interrogation of role that 5-HT_1B_ receptors might have on the developmental neurotoxicity of CPF.

Basal 5-HT_1B_ expression was markedly higher in females, compared to males. This is consistent with previous reports of higher basal levels of 5-HT synaptic proteins (5-HT_1A_ and 5-HT_2_) in females, compared to males (Slotkin and Seidler, 2005). For 5-HT_1B_ specifically, this sex difference is supported by previous work which reported differing functional consequences in the tail suspension and forced swim test following knockout of the 5-HT_1B_ receptor (Jones and Lucki, 2005). Reports of sex differences in 5-HT receptor expression in response to perinatal CPF exposure have been mixed, depending on the targeted brain region, but the effects has been generally greater in males than females (Aldridge et al., 2004; Slotkin and Seidler, 2005). However, we did not identify differences in the response to CPF exposure with our current model.

### Conclusions

The present study furthers our understanding of the underlying neurophysiological effects of gestational exposure to CPF. The enhanced spontaneous neuronal activity, increased expression of the receptor 5-HT_1B_, and the resulting dysregulation of PV neurons, are all potential causal mechanisms of the persistent neurobehavioral disorders associated with this exposure. Increasing our understanding of this complex phenomenon could eventually lead to preventative interventions and more informed decision making on the use of OP insecticides.

## Notes

**Acknowledgements:** JK is supported by the Department of Defense (DoD) Science, Mathematics, and Research for Transformation (SMART) Scholarship-for-Service Program. The content is solely the responsibility of the authors and does not necessarily represent the official views of the DoD.

### Competing Interest Statement

The authors have declared no competing interest.

